# mEthAE: an Explainable AutoEncoder for methylation data

**DOI:** 10.1101/2023.07.18.549496

**Authors:** Sonja Katz, Vitor A.P. Martins dos Santos, Edoardo Saccenti, Gennady V. Roshchupkin

## Abstract

In the quest to unravel the mysteries of our epigenetic landscape, researchers are continually challenged by the relationships among CpG sites. Traditional approaches are often limited by the immense complexity and high dimensionality of DNA methylation data. To address this problem, deep learning algorithms, such as autoencoders, are increasingly applied to capture the complex patterns and reduce dimensionality into latent space. In this pioneering study, we introduce an innovative chromosome-wise autoencoder, termed mEthAE, specifically designed for the interpretive reduction of methylation data. mEthAE achieves an impressive 400-fold reduction in data dimensions without compromising on reconstruction accuracy or predictive power in the latent space. In attempt to go beyond mere data compression, we developed a perturbation-based method for interpretation of latent dimensions. Through our approach we identified clusters of CpG sites that exhibit strong connections across all latent dimensions, which we refer to as ‘global CpGs’. Remarkably, these global CpGs are more frequently highlighted in epigenome-wide association studies (EWAS), suggesting our method’s ability to pinpoint biologically significant CpG sites. Our findings reveal a surprising lack of correlation patterns, or even physical proximity on the chromosome among these connected CpGs. This leads us to propose an intriguing hypothesis: our autoencoder may be detecting complex, long-range, non-linear interaction patterns among CpGs. These patterns, largely uncharacterised in current epigenetic research, hold the potential to shed new light on our understanding of epigenetics. In conclusion, this study not only showcases the power of autoencoders in untangling the complexities of epigenetic data but also opens up new avenues for understanding the hidden connections within CpGs.

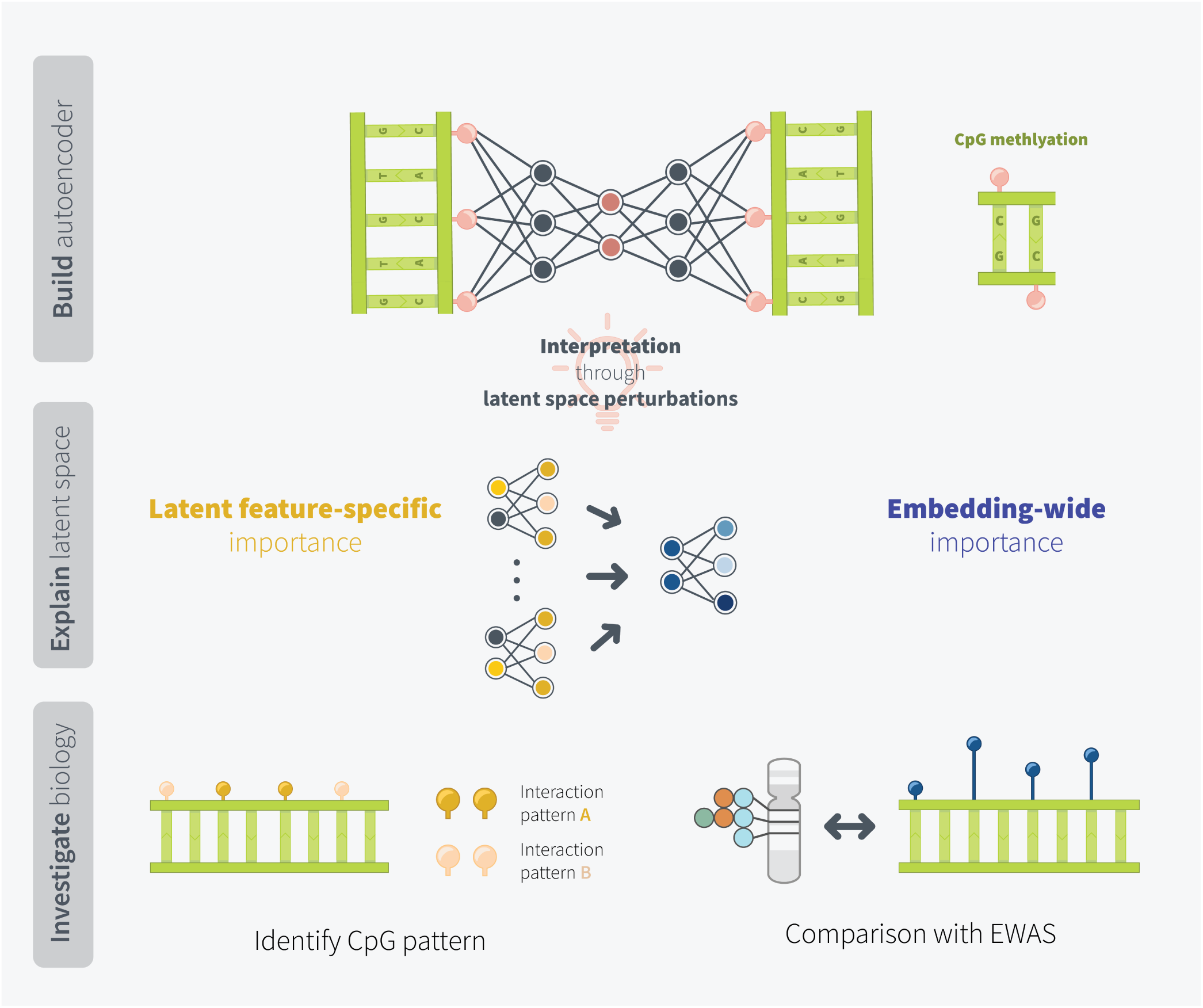

## 2 Introduction

Epigenetics comprises all changes in gene expression that are independent from DNA sequence alterations. DNA methylation, the most extensively studied epigenetic mechanism [1], involves the addition of a methylation group to a nucleotide, typically the cytosine residue of a cytosine-guanine (CpG) dinucleotide. Epigenome-wide association studies (EWAS) of high-throughput sequencing data have uncovered strong methylation signatures in response to extrinsic factors, such as environmental changes, as well as intrinsic factors such as stress [2–5]. Furthermore, the integrity of the methylome has been closely linked to healthy ageing, as aberrant DNA methylation patterns have been linked to age-related diseases, including Alzheimer disease, cardiovascular diseases, and cancer [6].

The reversible conversion between methylated (hypermethylated) and de-methylated (hypomethylated) CpG regions has been shown to be highly dynamic [7]. EWAS aim at examining the effect of epigenetic alterations on genome function by identifying DNA methylation variants statistically significant associated with phenotypes of interests. Using general linear models EWAS identify differentially methylated positions by conducting pairwise comparison of mean CpG methylation values between different phenotypes [8].

Despite the wealth of knowledge generated through EWAS [9], such univariate linear analyses are incapable of taking into account the spatial relationships of the up to 450.000 CpG sites measured simultaneously on each array [10]. However, as the genomic distribution of DNA methylation is tightly linked to its gene regulatory functions, gaining a deeper understanding of the relationship between CpG sites is crucial for comprehending how epigenetic modifications influence gene expression and thus ultimately have an impact on human health and diseases [11]. A recent study [12] demonstrated how ageing related CpGs can interact. They show that the contribution of individual CpGs to ageing can be fully dependent on secondary, interacting CpGs. In some instances, a biologically relevant primary CpG site can be completely silenced by a secondary hypermethylated CpG; upon hypomethylation of the respective secondary CpG site, the silencing effect disappears. We therefore argue that it is essential to expand the pool of methods capable of modelling high-dimensional methylomics data, with the goal of finding not only single CpGs of interest, but genome-wide methylation patterns.

One promising approach to overcome the limitations of EWAS in studying the relationship between CpGs is the application of autoencoders to genome-wide DNA methylation data. Autoencoders, and variations thereof, are versatile deep learning frameworks capable of non-linear dimensionality reduction [13–15], clustering [15], data generation and imputation [16–18], and performing classification and regression tasks [19, 20]. However, the application of autoencoders for whole genome methylation data has proven to be challenging due to their sensitivity to hyperparameter selection [21] and high computational cost [22]. Additionally, autoencoders, as other deep learning frameworks, are inherently not interpretable. Various approaches have been developed to approximate what an autoencoder learns. Most commonly, this involves visualisation of the latent dimension, revealing possible clusters or regions of interest [23–25]. While autoencoders are frequently being applied to DNA methylation data [13, 14, 19] little work has been conducted on interpreting individual latent features and exploring, for example, which CpGs share a relation through common latent features.

We hypothesise that the way an autoencoder groups together CpGs in its latent dimensions has biological meaning and might reveal novel insights regarding the relationship of CpGs. To explore this, we propose a chromosome-wise autoencoder framework for interpretable dimensionality reduction of methylation data (mEthAE). In an attempt to validate the performance of our autoencoder, we conduct supervised prediction of age and sex from the latent embedding. Our interpretability pipeline, which is based on latent feature perturbations, yields groups of related CpGs at the latent-feature specific (local), as well as embedding-wide (global) level, which allows us to study CpG relationships at different granularities. In an attempt to put the obtained local and global CpG groups into a biological context, we compare them to EWAS findings, assess their power to predict phentoypes (age), analyse their genomic location, associated biological pathways, correlation patterns, and genomic distances.

Bettering our understanding of the relationship between CpGs could, in the long term, contribute to our understanding of epigenetic mechanisms and have important implications for the development of novel therapeutic strategies for various diseases, as well as for the identification of biomarkers that can help diagnose and predict disease risk.

## 3 Results

### Architecture optimisation

To reduce the dimensions of genome-wide DNA methylation data we propose mEthAE, a framework based on chromosome-wise autoencoders. We decided to split our dataset per chromosomes and train 22 individual models, as the high number of input CpGs led to an explosion of trainable parameters if we attempted to train all chromosomes simultaneously. This parameter explosion disabled us from exploring using e.g. multiple hidden layers or larger latent dimensions. With 22 individual models, we were able to thoroughly explore and compare different hyperparameter settings (see material and methods). During optimisation, we focused on achieving a good trade-off between size of the embedding and performances, measured as validation loss and reconstruction accuracy which is defined as the Pearson correlation between input and reconstructed CpGs. An overview of the best models, their latent sizes, and reconstruction accuracies for each chromosome can be found in Table 1. Overall, we found it possible to compress the input information to a factor of up to 400, while simultaneously achieving reconstruction accuracies close to the maximal possible value (Fig 1). We expected the number of optimal latent dimensions to correlate with the size of the input. Interestingly however, although the number of CpGs for each chromosome greatly varies, we observed good performances with latent sizes of 70 across all chromosomes. During optimisation, we often observed a spike in performance around this number of latent dimensions, whereas performances only increased marginally when significantly more latent features were added (Supplementary Fig 2). To judge whether we encoded biologically meaningful information, we combined the latent dimensions of all autoencoders, totalling 1389 latent features, and conducted supervised prediction of age and sex. Age regression yielded a coefficient of determination of 0.82 and a RMSE of 8.72 years, while the classification AUC ROC was 0.95, clearly displaying the combined autoencoders captured biological signal in their embedding (Supplementary Fig 3).

**Figure 1:**
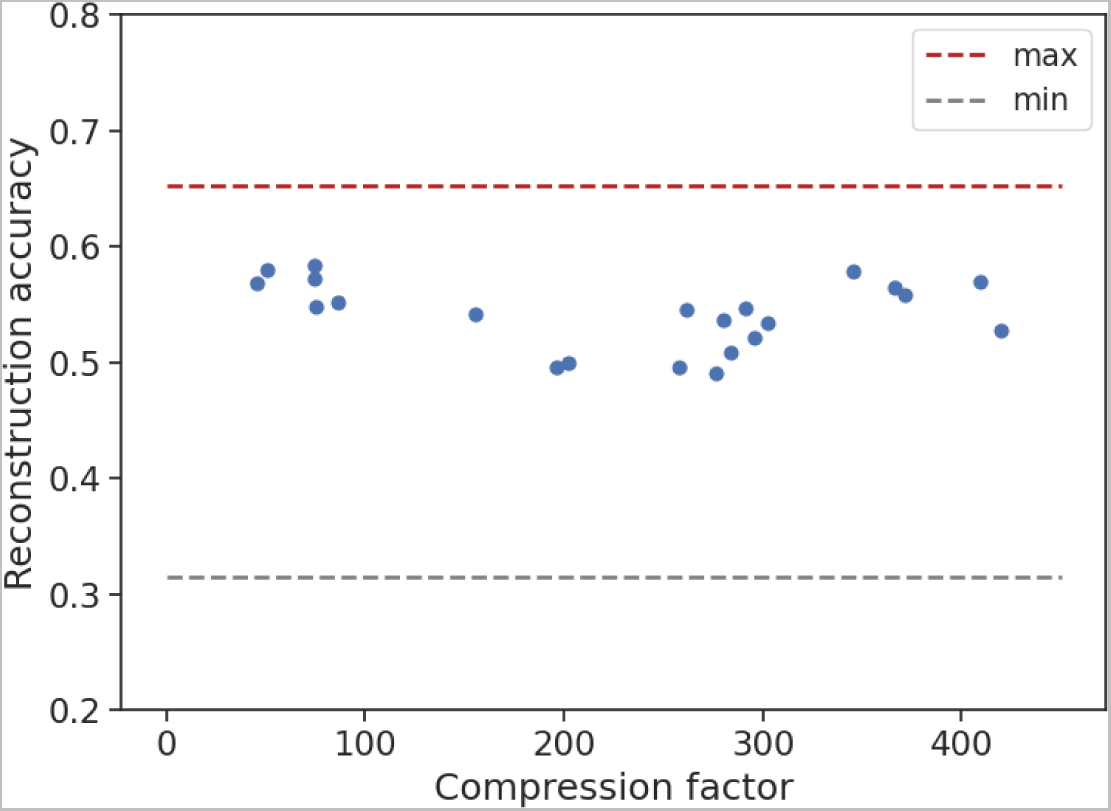
Comparison of reconstruction accuracies in relation to the compression factors for each chromosome. Red dashed line: reconstruction of models with a latent size equivalent to the number of input CpGs (no compression), averaged over all chromosomes; grey dashed line: reconstruction of models with a latent size of 1 (maximal compression), averaged over all chromosomes. Details on the models with no and maximum compression can be found in Supplementary Table 2.

**Table 1:** Overview of models after hyperparameter optimisation.

|  | Number of CpGs | Latent Size | Reduction Factor | Reconstruction accuracy |
| --- | --- | --- | --- | --- |
| **chr1** | 29482 | 72 | 409 | $0.56 \pm 0.21$ |
| **chr2** | 21984 | 60 | 366 | $0.58 \pm 0.21$ |
| **chr3** | 15547 | 45 | 345 | $0.58 \pm 0.21$ |
| **chr4** | 13100 | 50 | 262 | $0.55 \pm 0.21$ |
| **chr5** | 15965 | 38 | 420 | $0.53 \pm 0.22$ |
| **chr6** | 23070 | 78 | 296 | $0.52 \pm 0.21$ |
| **chr7** | 19599 | 70 | 280 | $0.54 \pm 0.21$ |
| **chr8** | 13704 | 88 | 156 | $0.54 \pm 0.22$ |
| **chr9** | 6619 | 88 | 75 | $0.55 \pm 0.22$ |
| **chr10** | 15736 | 52 | 303 | $0.53 \pm 0.21$ |
| **chr11** | 18570 | 50 | 371 | $0.56 \pm 0.21$ |
| **chr12** | 15165 | 52 | 292 | $0.55 \pm 0.21$ |
| **chr13** | 7802 | 90 | 87 | $0.55 \pm 0.22$ |
| **chr14** | 9417 | 48 | 196 | $0.50 \pm 0.23$ |
| **chr15** | 9715 | 48 | 202 | $0.50 \pm 0.22$ |
| **chr16** | 13823 | 50 | 276 | $0.49 \pm 0.22$ |
| **chr17** | 17044 | 60 | 284 | $0.51 \pm 0.23$ |
| **chr18** | 3654 | 72 | 51 | $0.58 \pm 0.22$ |
| **chr19** | 15475 | 60 | 258 | $0.50 \pm 0.22$ |
| **chr20** | 6567 | 88 | 75 | $0.58 \pm 0.21$ |
| **chr21** | 2719 | 60 | 45 | $0.57 \pm 0.19$ |
| **chr22** | 5243 | 70 | 75 | $0.57 \pm 0.21$ |
| | 300000 | 1389 | $233 \pm 121$ | $0.54 \pm 0.21$ |

Number of CpGs: number of input CpGs; Latent Size: number of nodes in latent embedding; Reduction Factor: fold reduction from input size compared to latent size; Reconstruction accuracy: Pearson correlation between input and reconstructed CpGs - indicated for each model by the mean and standard deviation reconstruction accuracy over all CpGs

### 3.1 Embedding-wide (global) CpG connectivity

By assessing the perturbation effects of CpGs across all latent features, we were able to estimate the embedding-wide (global) CpG relationships. We believe that the more impacted CpGs are by perturbations of the latent embedding, the stronger they are connected to (multiple) latent features. This degree of connectivity between latent features and CpGs - in short the CpG connectivity - reflects their importance in the network. CpGs exhibiting similar connectivities form what we refer to as global CpG connectivity groups. We sought to further characterise these global connectivity groups in terms of inter-cluster differences and potential biological relevance.

#### Degree of global connectivity reflects sample and probe heterogeneity

Our interpretability approach (see material and methods) categorised all input CpGs in three different connectivity groups, shown in Table 2. Whereas most CpGs show no or a low connectivity (none, 28.8%; low, 60.9%), a small percentage of CpGs were highly connected across the embedding (high, 10.3%). This ratio of highly connected CpGs was similar across all chromosomes, indicated by the low standard deviation, whereas the ratio of lowly and not connected CpGs differed substantially between chromosomes. In-depth comparison of these different global groups revealed clear differences between them. When inspecting the range of beta values, CpGs in the highly connected groups showcased a wide range of values, while CpGs in lower global connectivity clusters increasingly converge towards values of 0 or 1 (Figure 2A). The inter-sample heterogeneity, indicated through the standard deviation of beta values across samples, correlated directly with the degree of connectivity, with highly connected CpGs showing the highest standard deviations between samples (Figure 2B).

**Figure 2:**
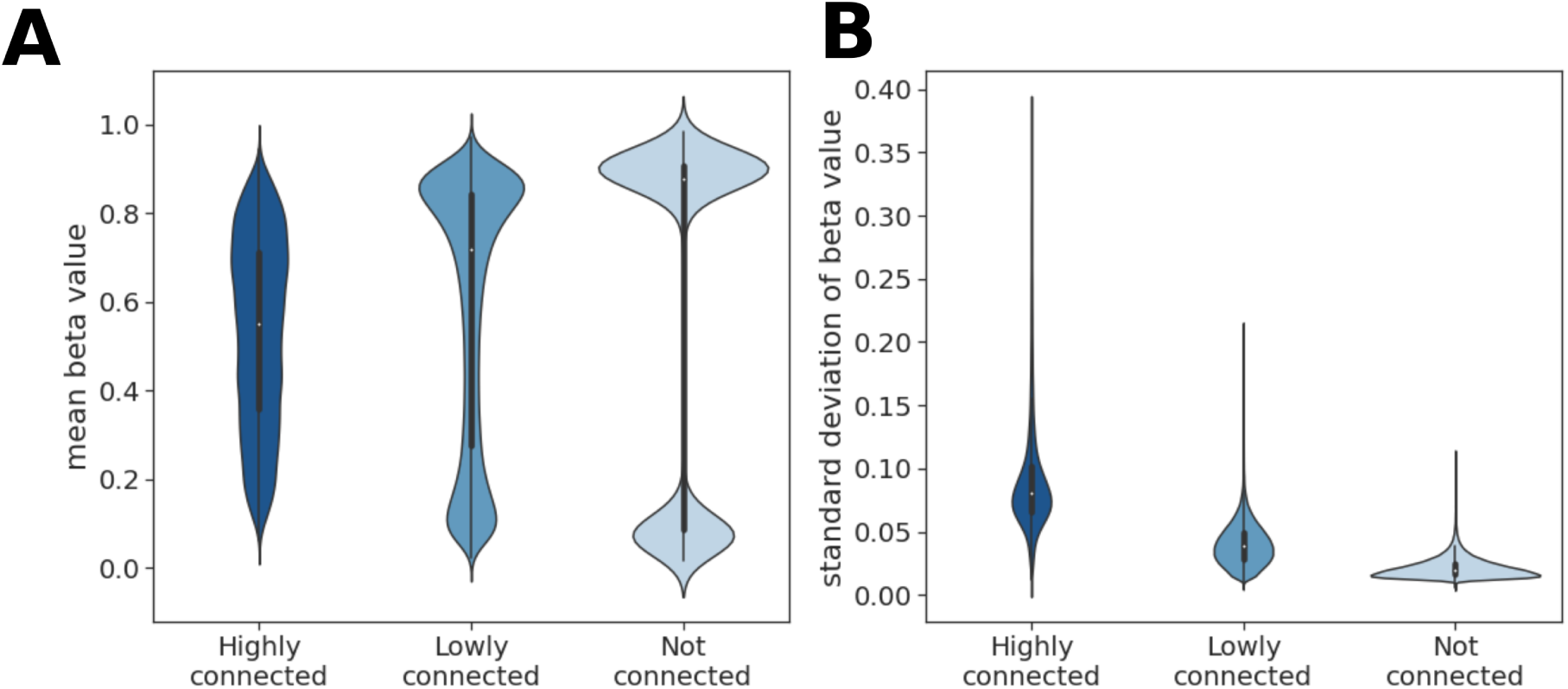
Characterisation of global connectivity groups. (A) average and (B) standard deviation of CpG beta values across all samples for the different clusters.

**Table 2:** Characteristics of global CpG groups.

| **Global connectivity group** | **Ratio of associated CpGs** | **Total number of CpGs across all chromosomes** |
| --- | --- | --- |
| global-high | $10.3 \pm 1.3\%$ | 30949 |
| global-low | $60.9 \pm 14.6\%$ | 181798 |
| global-none | $28.8 \pm 15.4\%$ | 87523 |

Normalised with regard to the total number of CpGs on each chromosome and average across all chromosomes

#### Globally highly connected CpGs show higher biological relevance

To estimate whether a distinction in biological relevance between CpGs from different global connectivity groups can be made we (I) interrogated how frequently CpGs from groups are reported in EWAS studies (Fig 3A) and (II) investigated their predictive power by assessing how accurately participants’ age can be predicted using CpGs from the different groups (Fig 3B). Comparison to EWAS studies revealed that CpGs from different groups are reported with differing frequencies (Fig 3A, dashed lines). To put this finding into relation, we employed a bootstrapping strategy (see Material and methods) (Fig 3A, densities). This combined approach revealed that highly connected CpGs are more often reported in EWAS studies than other groups or in the bootstrapped subset. The observed effect is still present, however less pronounced, with lowly connected CpGs and vanishes completely for not connected CpGs. We additionally conducted supervised age regression (Fig 3B, dashed lines) and evaluated in a similar bootstrapping approach as outlined above, namely through comparison against the prediction accuracies obtained with randomly drawn CpG subsets of the same size (Fig 3B, density). Similar to the picture obtained through the comparison to EWAS findings, it was observed that CpGs with high connectivities are more predictive than other clusters (bootstrap p-value *<* 0.05), while not connected CpG perform worse than a random subset. As a summary, using two independent approaches, we observed that the global connectivity of CpGs correlates with their biological relevance.

**Figure 3:**
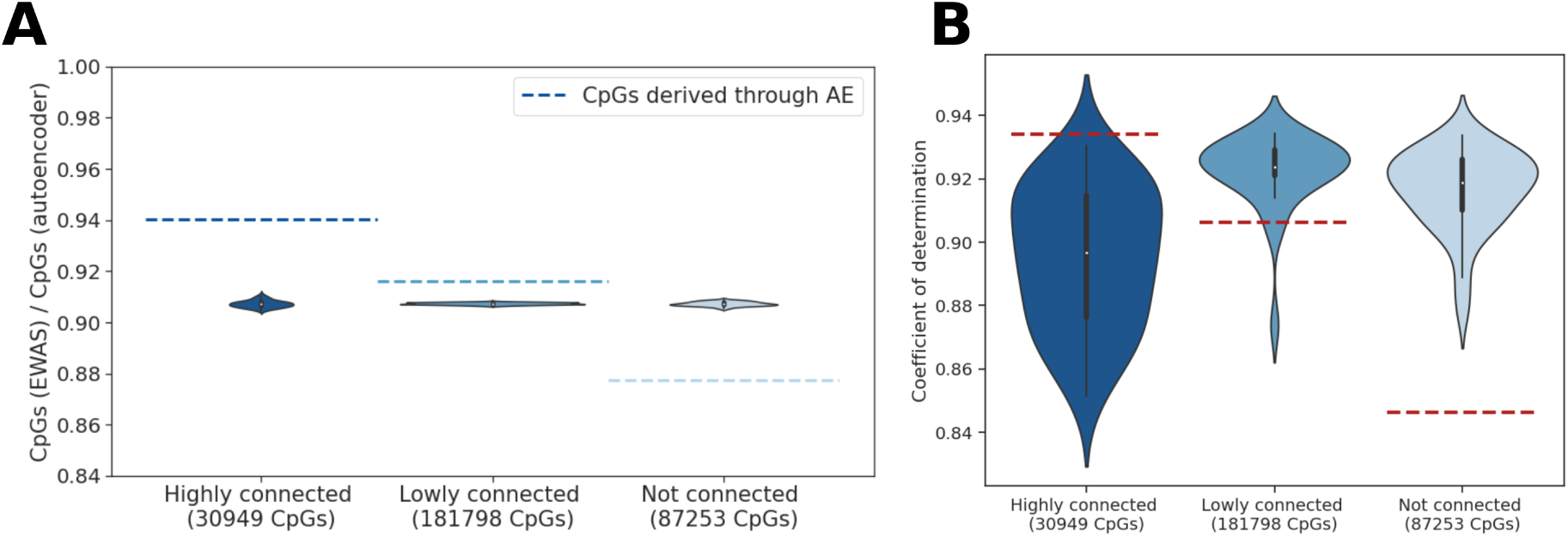
Analysis of biological relevance of globally highly connected CpGs. (A) Proportion of CpGs reported as significant in EWAS catalogue. (B) Supervised age prediction using CpGs from different global connectivity groups. Dashed line - CpGs recovered through interpretation approach; distribution - 100 times randomly sampled CpG set of the same size.

#### Latent feature-specific (local) CpG groups

In an attempt to gain a deeper understanding of the relationship between CpG sites, we focused on interpreting individual latent features, as we hypothesised that the way an autoencoder groups together CpGs in its latent dimensions has biological implications. By subsequently perturbing latent features and categorising CpGs into different groups based on the observed perturbation effects, we are able to delineate how strongly CpGs are linked to respective latent features. This approach yielded groups of high (local-high), medium (local-medium), low (local-low), and not (local-none) connected CpGs for each latent feature (see material and methods). An overview of the average number of CpGs assigned to each perturbation group can be found in Table 3. It is evident that a majority of CpGs are not affected upon perturbing a latent feature (89.4%); only a small percentage of CpGs show high (1.5%), medium (2.2%), or low (6.8%) effects. The e.g. local-high perturbation group only consists of around 208 *±* 107 CpG on average per latent feature.

**Table 3:** Proportion of CpGs in local perturbation groups.

| Perturbation group | Total number of CpGs<sup>1</sup> | Ratio of associated CpGs<sup>1</sup> |
| --- | --- | --- |
| local-high | $208 \pm 107$ | $1.5 \pm 0.2\%$ |
| local-medium | $296 \pm 144$ | $2.2 \pm 0.5\%$ |
| local-low | $930 \pm 475$ | $6.8 \pm 1.8\%$ |
| local-none | $12202 \pm 6127$ | $89.4 \pm 2.2\%$ |

<sup>1</sup>averaged across all latent features
averaged across all latent features and chromosomes

Following the observations from the analyses of global CpG groups, namely that globally highly connected CpGs seem to have biological implications, we aimed to investigate this further at a latent feature-specific level, hypothesising that CpGs part of the same local-high perturbation group are functionally or biologically connected. We thus tested whether these CpGs share genomic context associations, biological pathways, correlation patterns, or are spatially located in close vicinity to each other.

#### Latent feature-specific CpGs are enriched in specific genomic sites

Inspecting the genomic contexts annotated to the local-high CpGs in each latent feature proved that these CpGs are significantly more often located in genomic regions of interest (Table 4). Above 40% of latent features and their associated highly perturbed CpGs can be linked to Enhancer sites or 5‘UTR. Between 10-30% were enriched in DNase I Hypersensitivity Sites (DHS), the first exon, in the vicinity of the TSS (TSS1500), or at the 3‘UTR. It is to be noted that close to no association to CpG island, shelf or shore regions were found. The most common enriched site, with 62%, were gene bodies.

**Table 4:** Summary of highly perturbed CpGs for each latent feature significantly associated (p-value *<* 0.05) with different genetic contexts.

|  | Latent features associated<br>(p-value < 0.05) | Ratio [%] |
| --- | --- | --- |
| Gene body | 861 | 0.62 |
| Enhancer | 622 | 0.45 |
| 5'UTR | 586 | 0.42 |
| DHS | 383 | 0.28 |
| 1stExon | 305 | 0.22 |
| TSS1500 | 299 | 0.22 |
| 3'UTR | 223 | 0.16 |
| TSS200 | 55 | 0.04 |
| N-Shelf | 26 | 0.02 |
| S-Shelf | 19 | 0.01 |
| N-Shore | 17 | 0.01 |
| Island | 13 | 0.01 |
| S-Shore | 9 | 0.01 |

Ratio: Number of features normalised over the total number of latent features (n=1389)

#### Biological pathways, correlation patterns, or spatial proximity do not explain CpG grouping

Explanations behind the observed grouping of locally highly perturbed CpGs may include their biological roles, a similar co-correlation pattern, or proximity regarding their genetic locations.

To investigate a potential common biological role of CpGs in the local-high perturbation group, we tested for each latent feature if those CpGs are significantly associated with the same Gene Ontology (GO) term or Kyoto Encyclopedia of Genes and Genomes (KEGG) pathway (FDR *<* 0.05) (Supplementary Table 3). Out of 1389 latent features, only 201 showed association to (at least one) GO / KEGG term (14.5%). Notably, we observed big differences between chromosomes - while in most chromosomes not more than 15% of latent features could be linked to biological annotations, the CpG groups of every latent feature in chromosome 6 could be matched to a GO term (100%).

Deep learning models organise information following patterns in the data - we therefore examined possible co-correlation patterns between CpGs belonging to the same perturbation group by calculating their Spearman cross-correlation (Supplementary Fig 4). An average cross-correlation of *<*0.1 was observed, which suggests the absence of prominent correlations of monotonic associations.

Lastly, we hypothesised that CpGs are strongly encoded in a common latent feature due to spatial proximity on the chromosome, forming linkage disequilibrium (LD)-like clusters. To test this, we examined (I) the median pairwise distances of CpGs as well as (II) defined a LD-cutoff of 0.25 megabases (MB) and calculated a sparsity ratio for each latent feature, whereas a sparsity of close to 1 denote large spatial distances between CpGs (see material and methods). It is evident that CpGs highly perturbed for the same latent feature are spatially not clustered together on the chromosome, with median pairwise distances of *>*10 MB even for small chromosomes (Supplementary Fig 5 - A). This observation is confirmed upon inspecting the sparsity ratios for each chromosome (Fig 4). Even in the case of increasing the LD-cutoff, only a small decrease in the sparsity ratio is observed (Supplementary Fig 5 - B). An exception to this is chromosome 6 - it shows low median pairwise distances, a low sparsity ratio, as well as high sensitivity to variation in the LD-cutoff.

**Figure 4:**
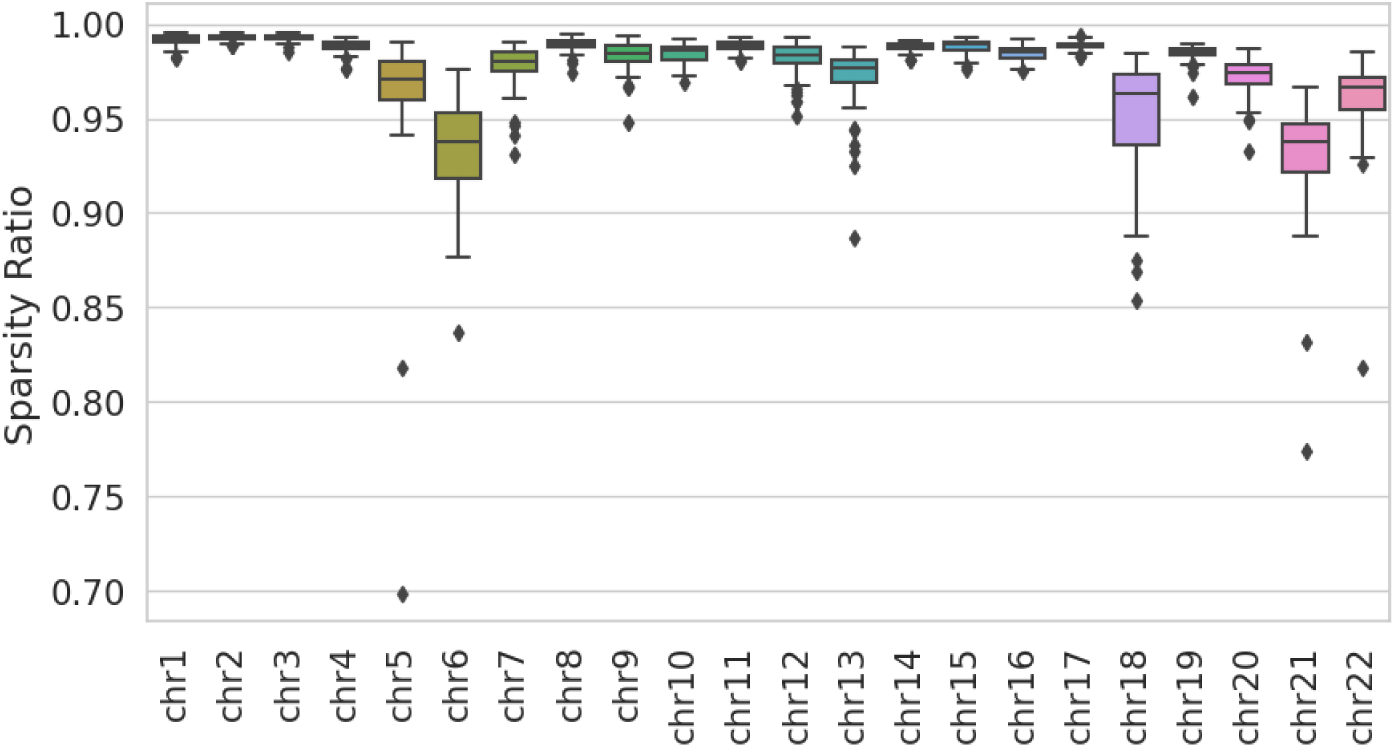
Sparsity ratio of local-high CpG perturbation clusters for all latent features with a fixed LD-cutoff of 0.25 MB. A sparsity ratio of 1 denotes distances of CpG sites above the designated cutoff, whereas 0 indicates identical positioning of CpGs.

## 4 Discussion

The application of deep learning on DNA methylation data is a promising approach having already made an impact in the field, but it remains challenging to delineate the learned biology from models. We therefore set out to build a framework not only capable of efficiently encoding methylation array data, but also enabling interpretation of how CpGs are grouped together in the latent embedding.

### High compression in latent embedding suggests signal sparsity

Although the Illumina 450K array only covers 2% of CpG sites of the human genome [26] and are therefore considered very sparse, our work reveals a high degree of redundancy in the data, demonstrated by the feasibility of compressing the input in a few latent features. We show that just 1389 latent features - as opposed to 300.000 input CpGs, which translates to compression of up to 400-fold - suffice to successfully achieve high reconstruction accuracies. Our results align with previously published deep learning models, which found comparable compression factors [13, 14, 19].

### Global CpG groups capture biological diversity

With the aim of interpreting which information is compressed in individual latent features, we developed an interpretability pipeline based on latent feature perturbations. Perturbing or permuting nodes of interest and observing the downstream effects on the network is a widely used attribution method for many data types [27–29]. A recent publication by Minoura et al. utilised the step-wise perturbation of hidden nodes to find the correlations of genes of interest with individual latent features, ultimately deducing regulatory programs associated with each latent dimension [30]. We took inspiration from these approaches and applied them to methylation array data. To facilitate a more structured interpretation we divided the obtained perturbations effects in two levels: a local, latent feature-specific level, as well as a global embedding-wide level, obtained by grouping CpGs that show perturbations across multiple latent features.

Our analysis of global CpG groups shows a clear pattern: the more these groups are influenced by perturbations of latent features, the stronger their connection to these features, and the more biological information they seem to carry. This becomes evident when we compare groups with varying levels of global connectivity based on their methylation beta values. Firstly, groups with higher connectivity usually contain CpGs with beta values around 0.5, a level often associated with significant variability in methylation patterns across different cells. Secondly, groups with higher connectivity also display a higher variability of beta values across samples, indicated by a high standard deviation. This is of interest, as typically probes with low variability (low standard deviation) across samples are deemed less biologically significant and are commonly removed during pre-processing. Furthermore, we find that CpGs with higher connectivity are more frequently reported in significant findings from epigenome-wide association studies (EWAS) and tend to show greater predictive power in supervised learning models. Together, this reinforces the idea that (i) the CpG groupings established by our autoencoder is biologically-driven, capturing meaningful relationships among CpGs of different chromosomes and (ii) we are able to observe this with our interpretability method, which suggests that CpGs in the high-global connectivity group may be key in capturing biological variations.

### Local groupings suggest potential long range non-linear CpG interactions

Building on the understanding that global CpG groups possess biological significance, our study ventured to explore the reasoning behind the local, latent-feature specific CpG groupings. Our investigations revealed that these CpGs often occupy crucial genomic sites like enhancers, exons, or gene bodies.

This is of particularly interest as it aligns with recent findings indicating the substantial impact CpGs located outside transcription start sites have on gene regulation [31, 32], a departure from earlier focus areas in research [33]. For example, enhanced methylation in gene bodies has been linked to increased gene expression [34, 35], while the methylation of first exons is closely associated with gene silencing [36]. Notably, enhancer methylation has recently emerged as a critical regulatory element, drawing significant attention in genomics [37–39].

Although this discovery resonates with our broader analysis, it doesn’t completely clarify the specific relationships within the CpG groupings. Our association of latent feature groupings with biological annotations like GO and KEGG terms indicates that only a fraction can be linked to biological processes or pathways. This suggests the absence of a uniform pattern across these groupings.

Additionally, we found low cross-correlation within these groups together with low ratio of short to mid-range CpG interactions. Instead, we observed that CpGs within the same perturbation group were dispersed along the chromosome, with considerable distances between adjacent sites, suggesting the presence of a long distance regulatory mechanism. A notable exception was chromosome 6, where we observed tighter clustering of CpGs within the same perturbation group, as evidenced by lower sparsity ratios and median pairwise distances. This aligns with the known genetic density and complexity of chromosome 6, which houses key immune response regulators like the human leukocyte antigen (HLA) complex [40, 41].

While our analyses suggest that the autoencoder groups CpGs based on long-range, non-linear interaction patterns, these patterns are not yet fully characterised in the current research landscape. While strong associations between nearby CpG sites are well known to impact gene expression [11, 42–44] the regulatory role of distal CpG sites is an underrepresented topic because of the difficulty in defining CpG-target gene relationships [45]. However, the importance of long distance CpG sites is not to be underestimated, as it has been shown that most genes are associated with *trans*-methylation sites more than 10 Mb from their promoter region, which explain on average 50% of the variation in gene expression [46]. Long distance CpGs have also been recently proven to be putative biomarkers for subtype-independent breast cancer diagnosis [45]. With the recent finding on the importance of ultra-long-range interactions between regulatory elements [47], we are now prompted to re-imagine the role of long distance methylation marks on gene modulation. However, the challenge to extract novel biological insights from complex data using deep learning remains a daunting task [14].

Our study thus contributes to a deeper understanding of the genomic distribution of DNA methylation and its biological implications, marking a significant step forward in epigenetic research.

### Limitations

Due to the computational restrictions we trained individual autoencoder models on each chromosome. Therefore, our current framework can not detect cross-chromosomal interactions of CpGs. While this topic is underrepresented in literature, given our current findings it would be important to test such hypothesis and train one large model for all chromosomes [48, 49].

## 5 Conclusion

In this research, we have successfully developed chromosome-specific autoencoders for methylation array data, achieving a remarkable up to 400-fold reduction in data size without compromising the performance of our models. Our innovative approach, which includes a custom perturbation-based method, allowed us to explore both broad and specific CpG connections within the embedding.

Our finding that we are able to reduce the data size by 400-fold encourages a re-evaluation of thresholds applied in EWAS studies; the redundancies observed in the data suggest that the number of true independent tests may be lower than currently assumed for most studies, which would affect the EWAS significance threshold. A key finding from our interpretability analysis is that CpGs with strong connections across the entire embedding demonstrate significant biological relevance, corroborating findings from epigenome-wide association studies (EWAS). When examining CpGs grouped in individual latent features, we observed that these groupings did not correspond to typical correlation patterns or proximal locations on chromosomes.

This intriguing result points towards the possibility of long-range, non-linear interactions between CpGs, a phenomenon that has not been extensively characterised in epigenomic research. Our study thus opens up new avenues for understanding the complexities of DNA methylation and its role in gene regulation, indicating a rich field of study for future research in epigenomics.

## 6 Methods

### 6.1 Datasets

The data used derived from a continuous ageing study, publicly obtainable from the Gene Expression Omnibus (GEO) under accession number GSE87571. This study by Johansson et al. assessed the methylome of the white blood cells of 732 participants, ranging in age from 14 to 94 years old, using the Illumina Infinium 450K Human Methylation Beadchip [50]. For comparison to the EWAS studies, all metadata and results of the EWAS catalogue [51] were downloaded (version 2022-8-18). For analyses, only studies with results from measuring whole blood samples were included.

### 6.2 Preprocessing

We preprocessed the raw data using the *PyMethylProcess* pipeline, following the steps described in the original *PyMethylProcess* publication [52]. This yielded a total of 300.000 DNA methylation values, called beta values, which are continuous variables between 0 and 1, representing the ratio of methylated versus unmethylated probes for the same locus.

### 6.3 Computational framework

#### 6.3.1 Autoencoder

To reduce the dimensions of the DNA methylation data, our study utilised an autoencoder network, which is an unsupervised neural network consisting of an encoder and a decoder part (see Figure 5). While the encoder maps the high dimensional input data into a lower dimensional latent embedding, the decoder attempts to reconstruct the original input from said embedding. We trained the network through a mean squared error (MSE) loss function which quantifies the error between the original input and reconstruction. Prior to training models, the data set was divided into 70% training, 20% testing, and 10% validation.

**Figure 5:**
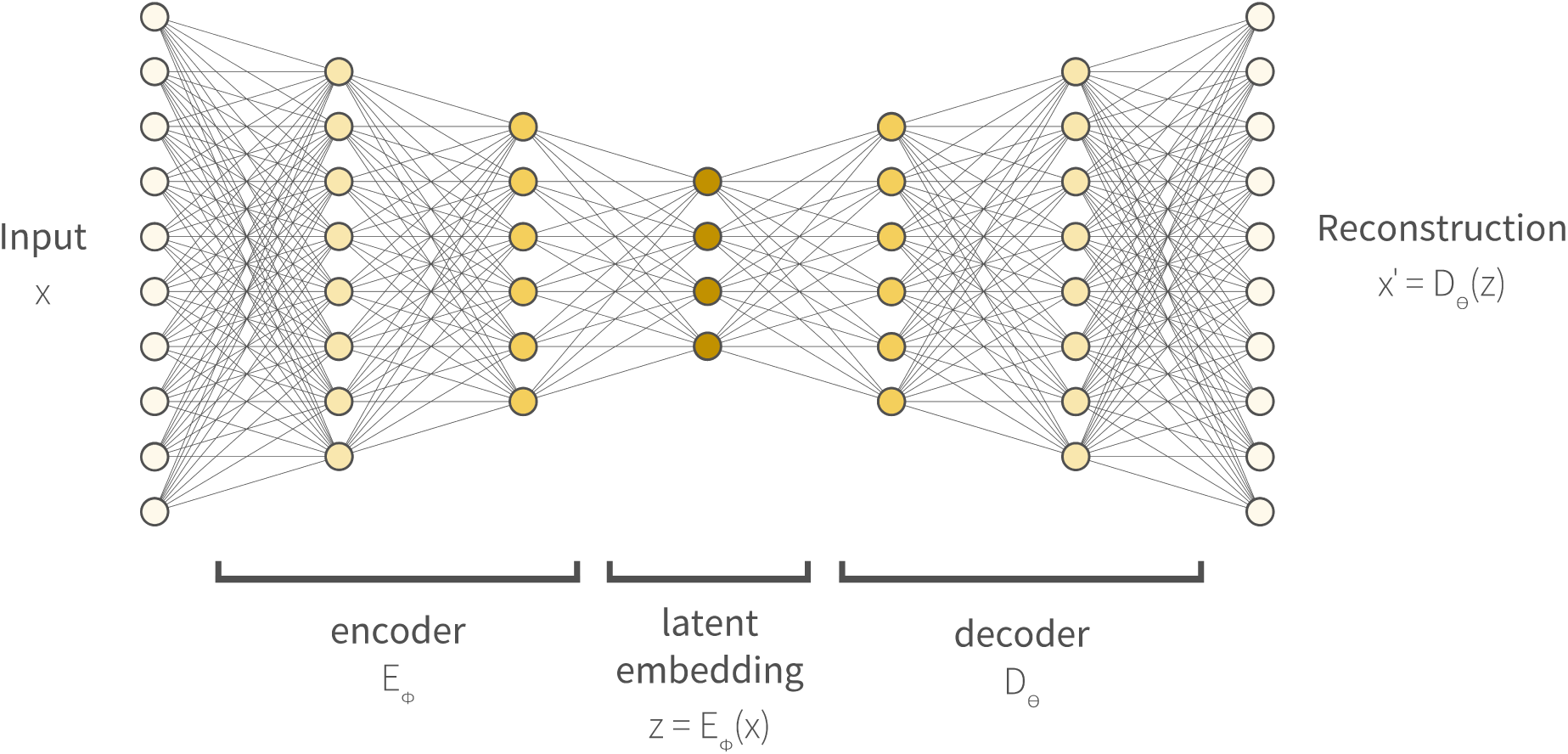
Schematic representation of an autoencoder. The input (*x*) denotes a set of input CpGs e.g. 29482 CpGs for chromosome 1. The encoder (*E_ϕ_*, parameterised by *ϕ*) consists of contiguous hidden layers, each with fewer nodes. The latent embedding (*z*) represents the bottleneck of the autoencoder with the minimum number of nodes. The decoder (*D_θ_*, parameterised by *θ*) reversely mirrors the layer structure of the encoder, with the final layer featuring the same number of nodes as the input layer as it attempts to reconstruct the original input from the latent embedding.

### Architecture optimization

To ensure that we do not compromise model design due to memory issues, we split our dataset chromosome-wise, ultimately training and optimising 22 separate models, one for each chromosome. To derive an optimal architecture for our autoencoders, we carried out an exhaustive hyperparameter search assessing the combination of multiple activation functions, layers (batch normalisation, dropout), learning rates, and number of latent features. All tested architectures were based on densely connected autoencoders with 2 hidden layers before the bottleneck (see Figure 5). Between each hidden layer, reduction factors of 3 and 2 were selected, resulting in a total reduction to 6% of the original input size respectively (Supplementary Table 1). Our hyperparameter optimisation strategy comprised two consecutive parts: firstly, a coarse scan to evaluate a broad range of hyperparameters, followed by a fine scan further optimising the best performing settings.

#### Coarse hyperparameter scan

During the coarse scan we evaluated latent space sizes with a reduction factor of 2, 4, 8, 16, 32, or 64 respective to the last hidden layer. As a regularisation method to avoid over-fitting, dropout layers with probabilities of 0.1, 0.3, and 0.5 were evaluated. Batch normalisation layers [53] were added to the encoder, which was observed to also enhance training speed and stability. Learning rates evaluated ranged from 0.0001 until 0.005. For the activation function, we used the Parametric Rectified Linear Activation Function (PReLU) [54] in all layers except the output layer, which employed a sigmoid activation function, as beta values are finite continuous values between 0 and 1. Models were trained for a fixed period of 500 epochs using an Adam optimiser [55] and a batch size of 64. As loss function mean squared error (MSE) was used. To evaluate and rank the training performance of models the loss on the validation set was monitored as well as the reconstruction accuracies, measured as the Pearson correlation coefficient (*ρ*) between the original and reconstructed CpGs. To identify an optimal latent space size from the coarse scan, we evaluated and ranked all combinations of hyperparameters for each latent space size. Subsequently, the latent size showing a satisfactory trade-off between validation loss, reconstruction accuracy, and number of dimensions was selected for further refinement in the fine hyperparameter scan.

#### Fine hyperparameter scan

To design the latent size grid for the fine hyperparameter scan, six latent sizes in close vicinity of the optimal latent space size from the coarse scan were selected. An overview of the final models including hyperparameters can be found in Supplementary Table 1. Subsequently the fine search followed the same strategy as the coarse search, assessing the identical set of hyperparameter combinations. Ultimately, our optimisation procedure yielded one hyperparameter-tuned autoencoder model per chromosome.

### Evaluation of reconstruction accuracies

To objectively evaluate whether the reconstruction accuracies (measured by the Pearson correlation (*ρ*) between the input and reconstructed CpGs) of our optimised models are satisfactory, we wanted to compare them to a version of the model without any compression in the latent space (number of latent dimensions = number of input CpGs), as well a version with maximum compression (number of latent dimensions = 1). In theory, the model without any compression should achieve the maximum possible reconstruction accuracy, while the model with only a single latent dimension denotes the minimum achievable accuracy. All models were built in *pytorch* (version 1.10.2) [56] and trained on using the GPU units RTX 2080 Ti (11GB).

#### 6.3.2 Supervised prediction of age and sex

The amount of biological information encoded by the autoencoders was estimated by trying to predict participants’ age and sex from the latent embeddings using a supervised model. Therefore, we used the *scikit-learn* [57] implementation of the Random Forest Regressor and Random Forest Classifier. Hyperparameters of the model were left unchanged, except for the number of trees in the forest, which was raised to 1000. Participants with missing annotation were excluded from the supervised prediction. To avoid data leakage, we used the same train-test splits for training the supervised model as were used for training the autoencoder. Scoring for age regression was done by calculating the coefficient of determination (*R*^2^) from actual and predicted age and for sex using the area under the ROC curve (AUC-ROC).

#### 6.3.3 Interpretability of latent features

To understand how the autoencoder encodes input information (CpGs) in the latent embedding, we implemented a post-hoc interpretation approach based on sequential perturbation of latent features which is outlined in the following sections. A graphical summary of the steps is provided in Figure 6.

**Figure 6:**
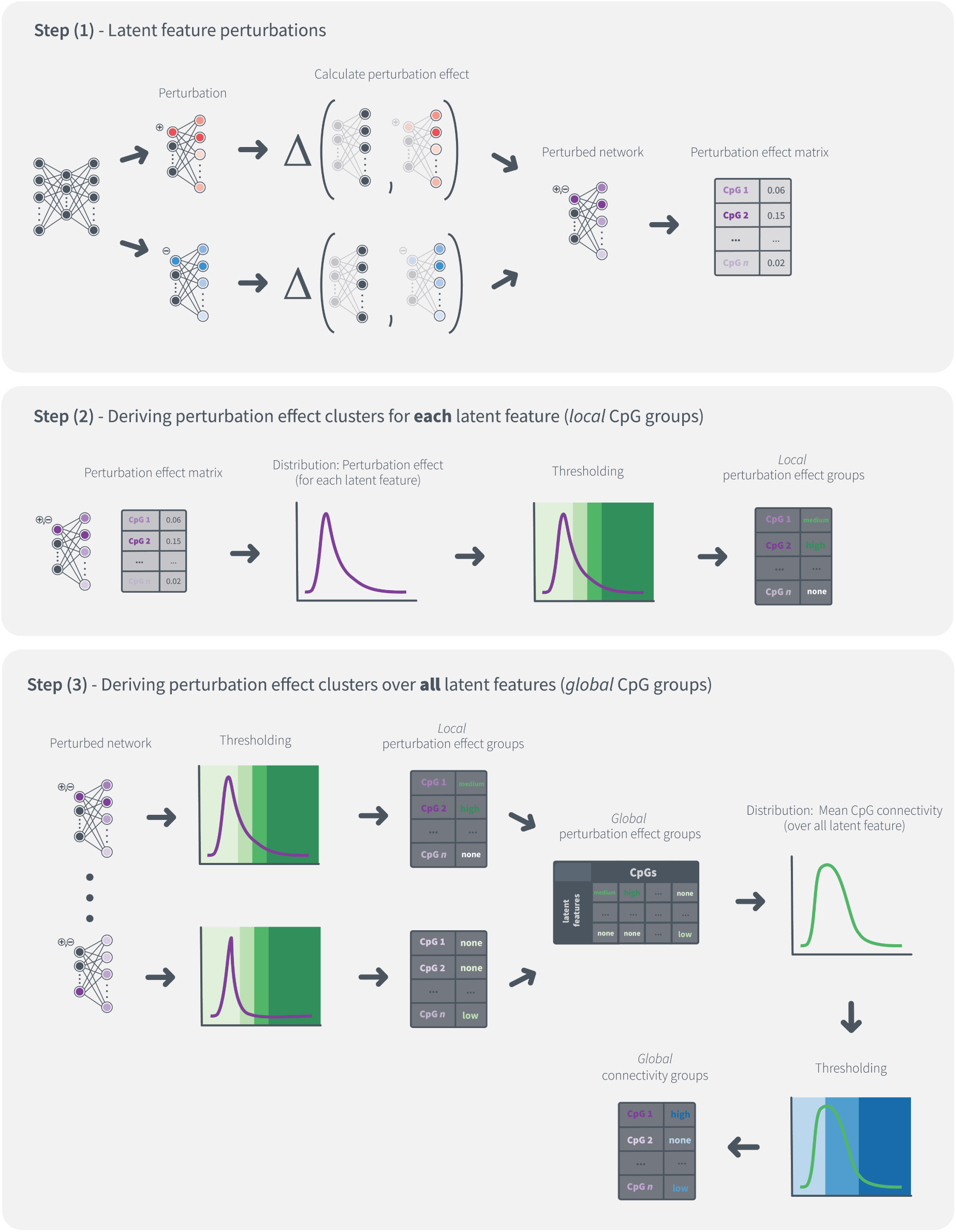
Graphical summary of the interpretability approach utilised in this work which bases on latent features perturbation. An in-depth description of each step can be found in section 6.3.3. An example of the observed perturbation effects can be found in Supplementary Figure 1.

### Step (1) - Latent feature perturbations

Latent feature perturbations were carried out sequentially one at a time while holding the remaining latent dimensions fixed. The perturbation value was determined to be one standard deviation of the latent feature activation. Subsequently, other forward passes were conducted, this time with the latent feature activation being altered by the addition or subtraction of the perturbation value. To estimate the effect of the introduced perturbation on each CpG, the median absolute difference between the original, unperturbed reconstruction and the new, perturbed reconstruction was calculated. Finally, the differences over all samples and both perturbation values were averaged, resulting in a single perturbation effect value for each CpG. Repeating this procedure for each latent feature yielded a perturbation matrix with a single perturbation effect value for each CpG and latent feature.

### Step (2) - Deriving perturbation effect clusters for each latent feature (local CpG groups)

For each latent feature we sought to categorise CpGs into different groups based on the perturbation effects ranging from highly, medium, lowly, to not perturbed groups - we refer to those as “local” CpG groups. To categorise CpGs into the aforementioned effect groups, we determined each latent feature’s mean perturbation effect and standard deviation and assigned latent-feature specific thresholds (Table 5). Subsequently, for each latent feature the perturbation effect of every CpG was examined and CpGs assigned to an effect group using these thresholds. It is important to highlight that CpG assignments were not unique, as these group assignments were performed for each latent feature sequentially.

**Table 5:** Latent-feature specific thresholds for assignment of CpGs to perturbation effect groups. Thresholds were derived through determining each latent feature’s mean perturbation effect and its standard deviation.

| perturbation effect group | threshold |  |
| --- | --- | --- |
|  | lower | upper |
| local-high | $> 3 * std()$ | - |
| local-medium | $> 2 * std()$ | $< 3 * std()$ |
| local-low | $> 1 * std()$ | $< 2 * std()$ |
| local-none | - | $< 1 * std()$ |

### Step (3) - Deriving embedding-wide perturbation effect groups (global CpG groups)

In addition to the latent feature-wise CpG grouping described above, we also sought to infer CpG groups based on all latent feature perturbations combined. We therefore used the effect groups derived from Step (2) to assess the “global” CpG relationships, which are the perturbation effects of individual CpGs across all latent features. Similar to the thresholding procedure described in Step (2), the mean perturbation effect of all CpG was derived and each CpGs classified into three global relationship groups, termed global CpG connectivitiy groups: globally highly, lowly, and not connected (Table 6).

**Table 6:**
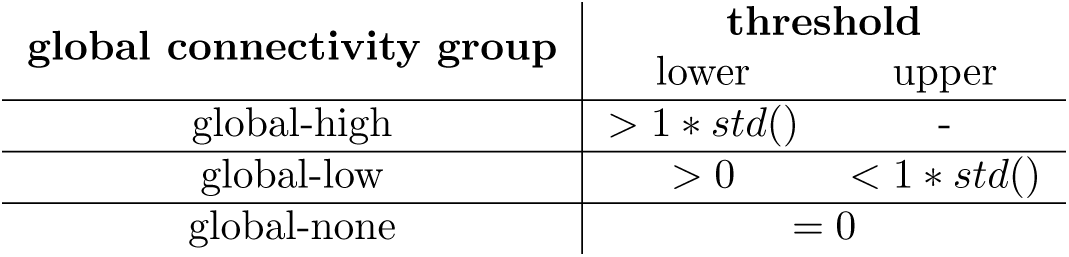
Thresholds for assigning CpGs to global connectivity groups.

| global connectivity group | threshold |  |
| --- | --- | --- |
|  | lower | upper |
| global-high | $> 1 * std()$ | - |
| global-low | $> 0$ | $< 1 * std()$ |
| global-none | $= 0$ | |

As a summary, our interpretability approach yields insights on two levels. Firstly, a detailed insight of which CpGs share a relationship by being linked to common individual latent features. Secondly, an estimate of how strongly CpGs are perturbed across the whole embedding, indicating their overall (global) connectivity in the network.

### 6.4 Characterisation of CpG clusters

#### 6.4.1 Comparison to EWAS findings

A biological validation of the global CpG groups derived by our interpretability pipeline was done through comparison to the EWAS catalogue, calculating how often CpGs from each connectivity group are reported in the EWAS catalogue. To estimate whether the CpGs found by our interpretability method were significantly over- or under-represented with regards to CpGs reported through EWAS, we followed the bootstrapping approach outlined here: (i) all CpGs of the perturbation group under study were checked for significant associations in EWAS studies and the number noted; (ii) a random set of CpGs of the same size as the CpG group under study was drawn and checked for significant associations in EWAS studies; (iii) repeat (ii) for 100 iterations to derive a distribution indicating the chance of a random group of CpGs being mentioned in EWAS; (iv) derive a significance measure by comparing the number of random CpG group configurations that showed the same or higher overlap with EWAS findings than our group of interest.

#### 6.4.2 Investigation of common genomic context, enrichment in biological pathways, correlation, and spatial location

We tested for potential common genetic location or functions of CpGs that are part of the same perturbation group by extracting the genetic information provided in the Illumina Infinium Methylation450K manifest (version 1.2). We applied hypergeometric testing to assess whether CpGs have a common location relative to the nearest gene, association to the same CpG island, location relative to the CpG island, regulatory feature group, DNase I Hypersensitivity Site (DHS), or are located in the same enhancer site. Additionally, using the missMethyl package [58] in R, we tested whether CpGs of the same perturbation groups belong to the same Gene Ontology (GO) terms or Kyoto Encyclopedia of Genes and Genomes (KEGG) pathways (significance threshold FDR *<* 0.05). To test for intra group correlation we examined the average Spearman cross-correlation of CpGs in the same perturbation group. To investigate whether CpGs connected to the same perturbation group form LD-like clusters on the chromosome, we assessed their spatial distances by calculating a CpG sparsity ratio. We therefore calculated the pairwise distances of CpGs within the same perturbation group and defined a LD-cutoff of 0.25 MB. The sparsity ratio was subsequently calculated by assessing the number of pairwise distances not within the LD-cutoff divided by the total number of distances.

### 6.5 Code availability statement

All code for framework development and performance analysis associated with the current submission is available at github.com/sonjakatz/methAE_explainableAE_methylation. Any updates will also be published on github. Preprocessed data as well as trained models are available upon request.

## Acknowledgments

The authors thank the supporters of this study, namely the European Union’s Horizon 2020 research and innovation programme under the Marie Sklodowska-Curie grant agreement No. 860895 Tran-SYS. Furthermore we thank the members of the Computational Population Biology group at Erasmus Medical Center for their critical and creative input to this work.

## Supplementary Material

**Supplementary Figure 1:**
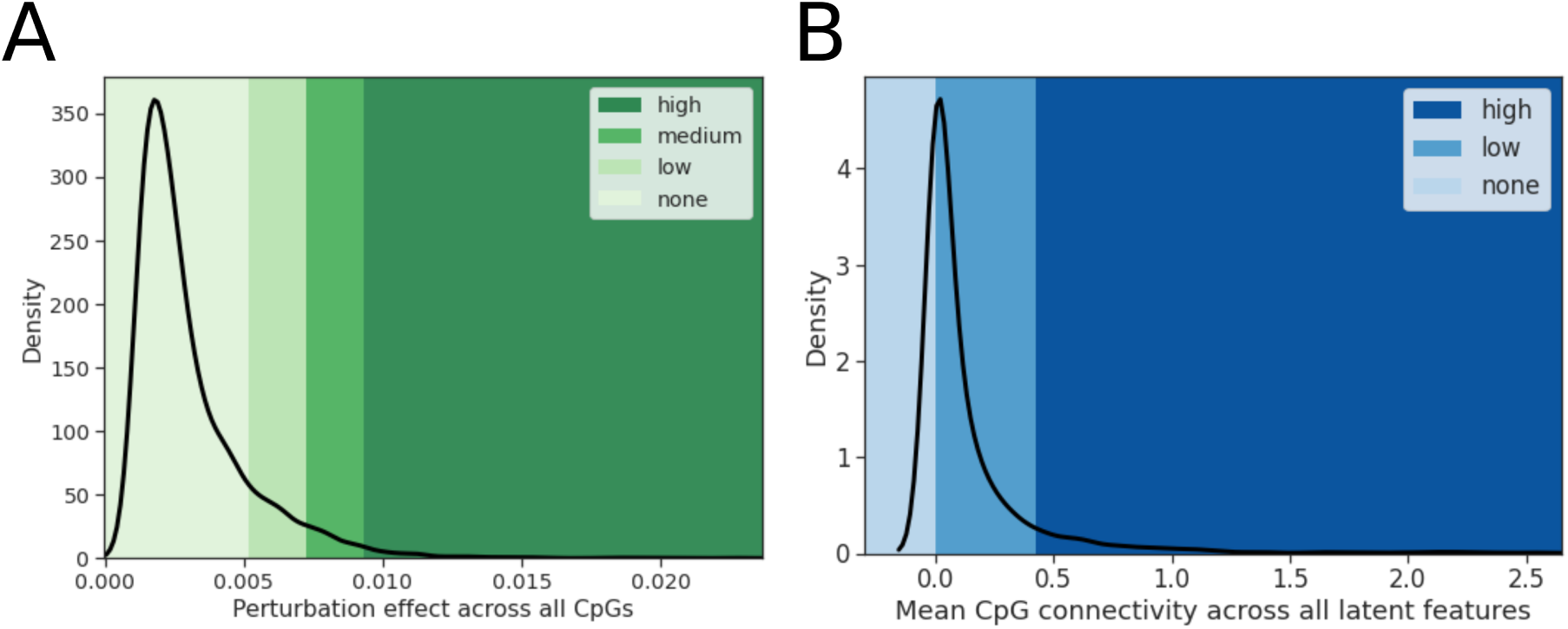
Example of observed perturbation effects using chromosome 22. (A) (Local) perturbation effect across all CpGs for a random latent feature. Colours indicate the applied thresholds for cluster assignment, for details see Table 1 in main manuscript. (B) (Global) CpG connectivities across all latent features. Colours indicate the applied thresholds for cluster assignment, for details see Table 2 in main manuscript.

**Supplementary Figure 2:**
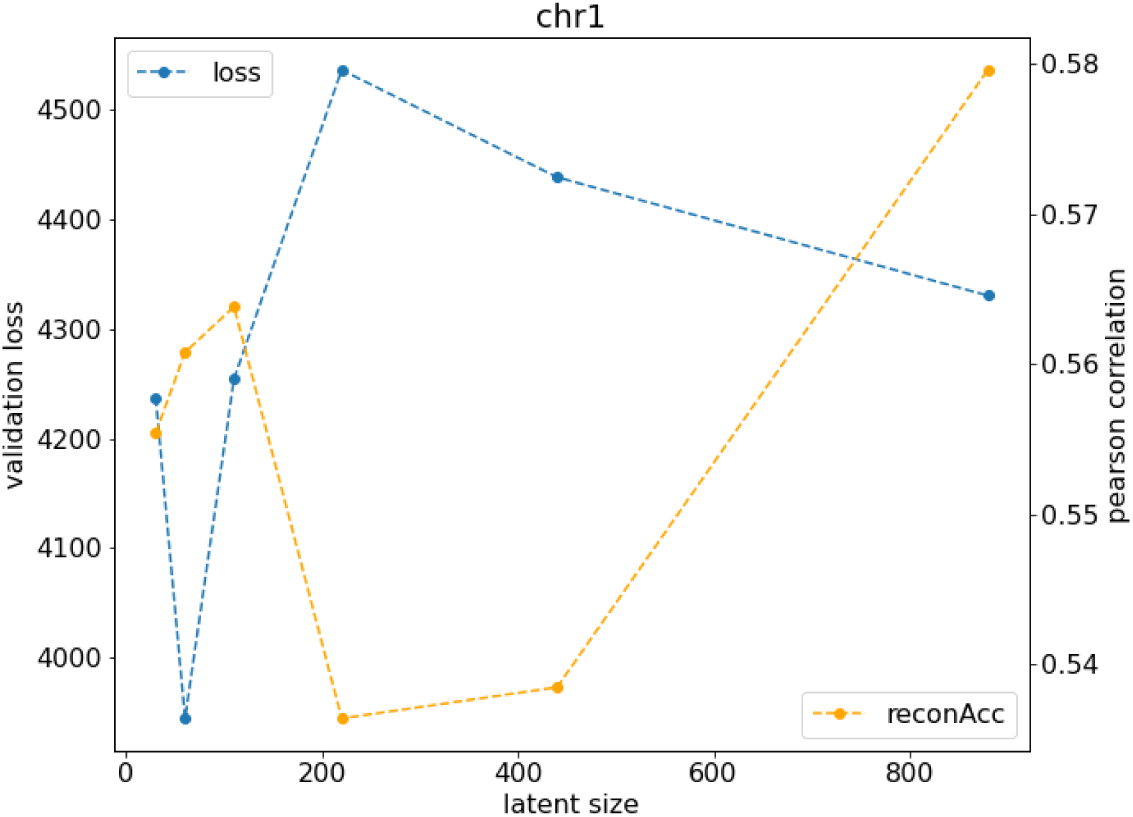
Rough hyperparameter scan results of chromosome 1. Results exemplary for other chromosomes; validation loss measured as mean squared error (MSE) loss between input and reconstruction; reconstruction accuracy measured as Pearson correlation between input and reconstruction, averaged over all CpGs

**Supplementary Figure 3:**
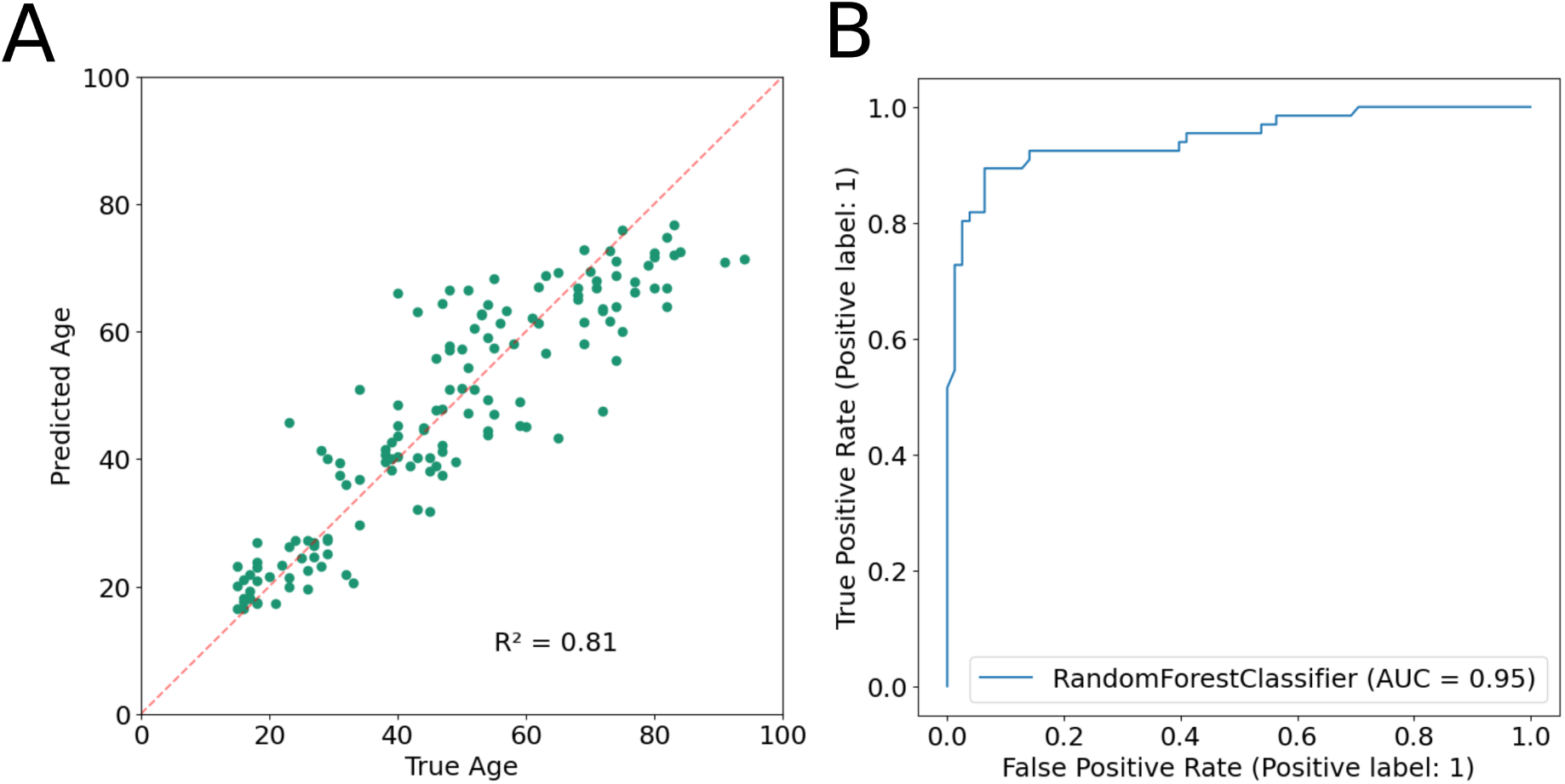
Supervised predictions using the combined latent embeddings of all chromosomes. (A) age regression, *R*^2^: coefficient of determination; (B) AUC-ROC curve of classification of participants’ sex, AUC: Area under the curve.

**Supplementary Figure 4:**
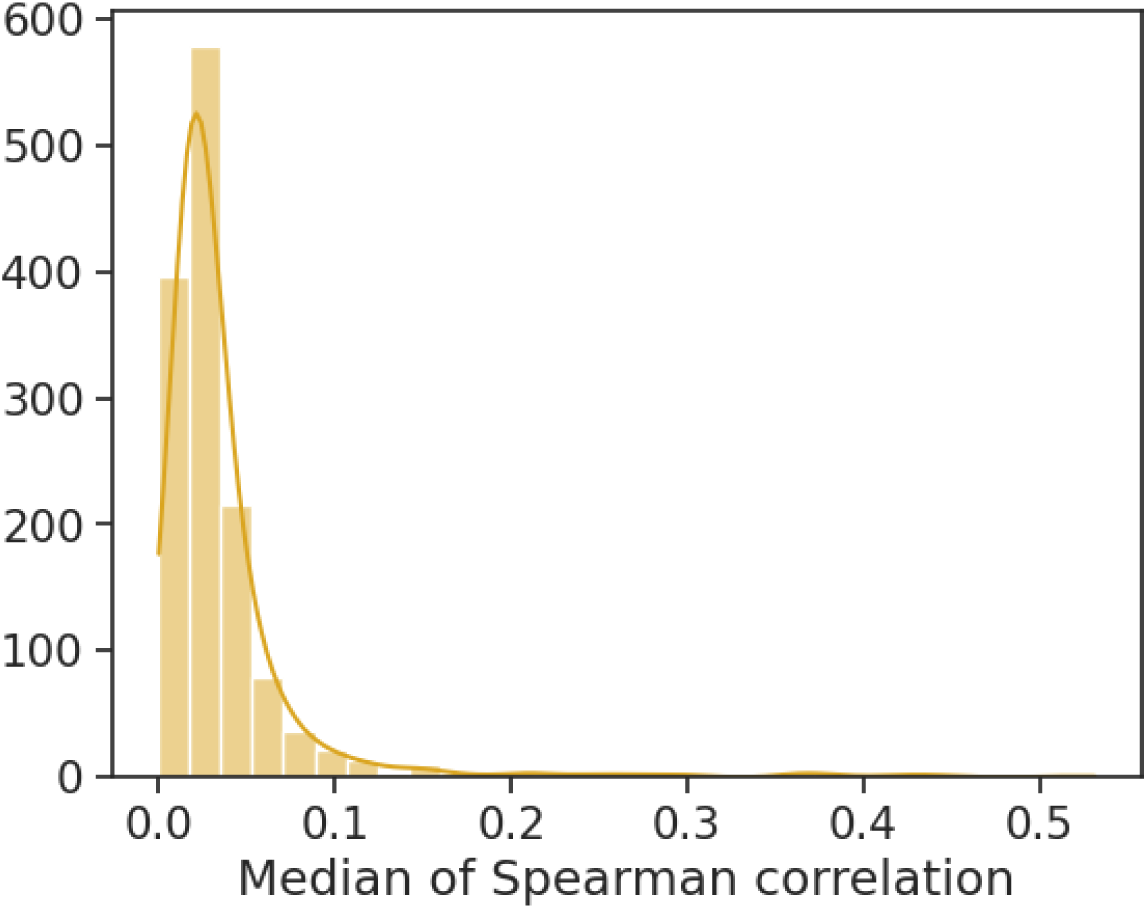
Median spearman cross-correlation over all latent features (n=1389).

**Supplementary Figure 5:**
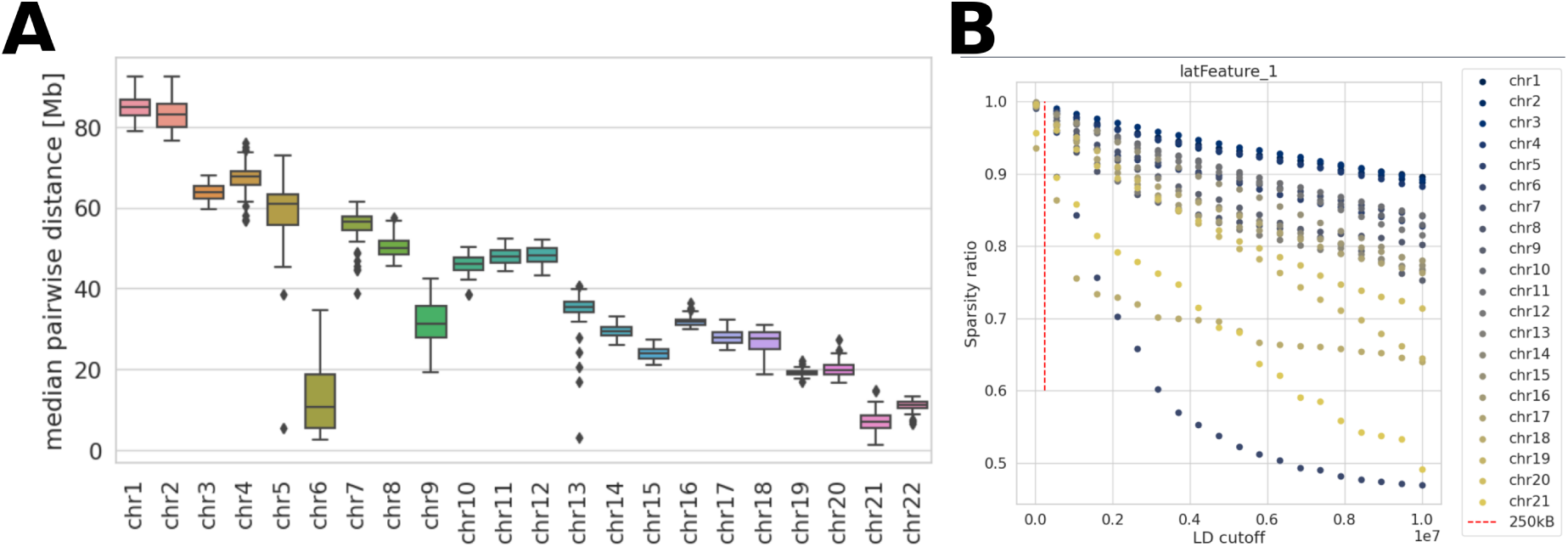
Median pairwise distances of locally highly connected CpGs. A) Median pairwise distances between local-high CpGs of the same latent feature; all latent features per chromosome shown. (B) Effect of the variation of LD-cutoff values on the sparsity ratio for each chromosome. One latent feature is shown as representative for all latent features.

**Supplementary Table 1:**
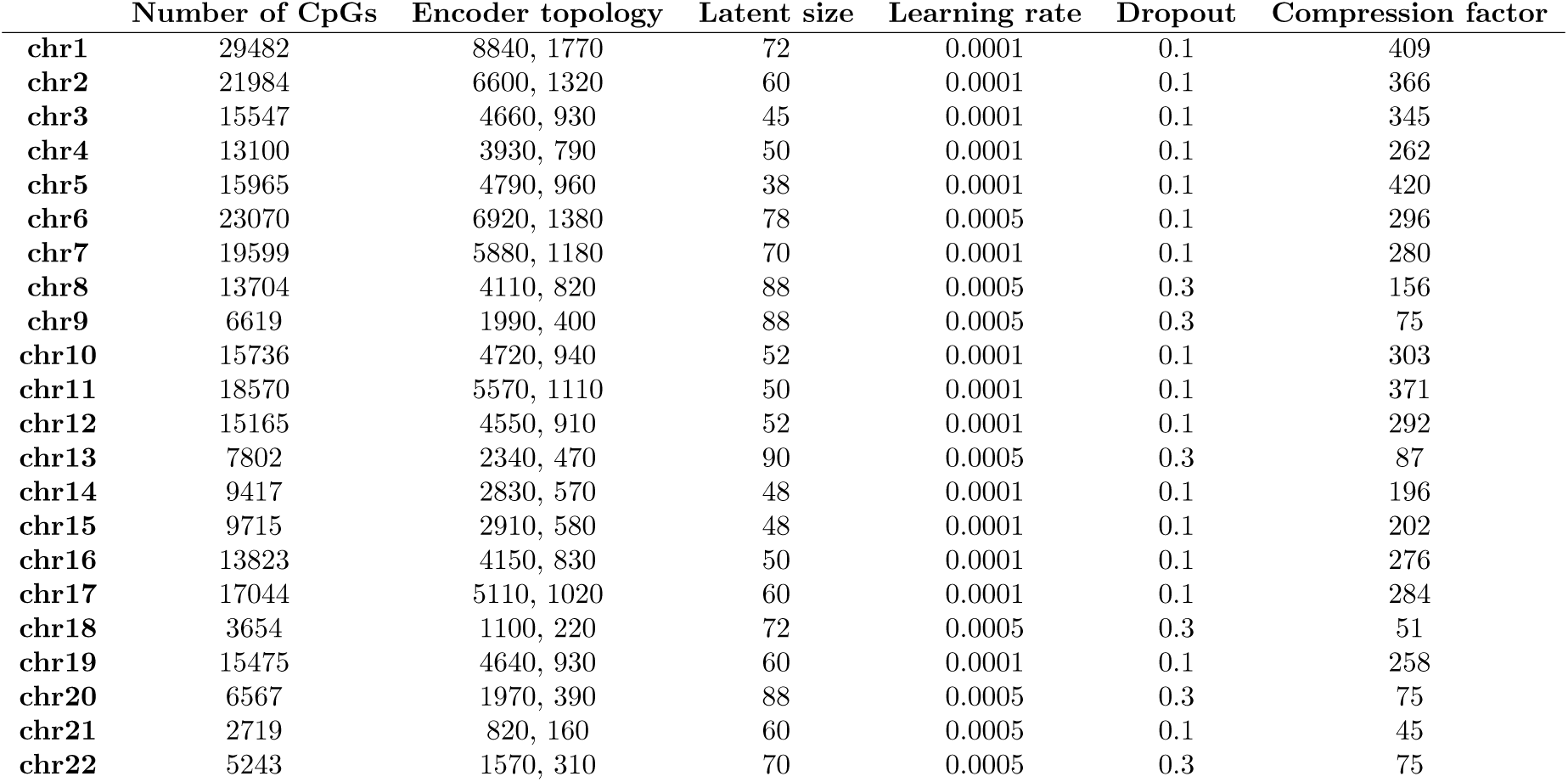
Details on architecture of all chromosome-wise autoencoders after hyperparameter optimisation.

**Supplementary Table 2:**
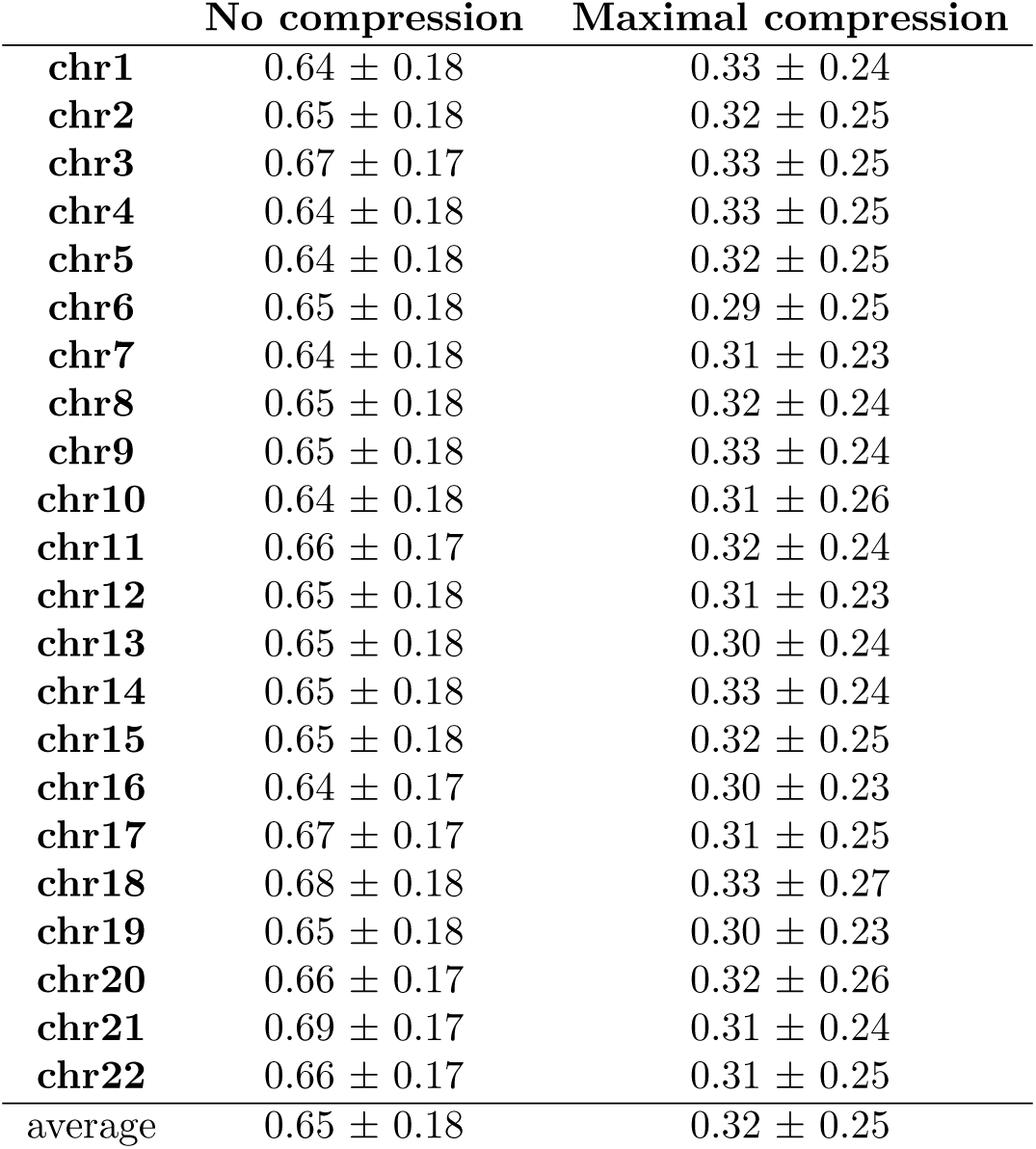
Overview of reconstruction accuracies for models with minimal and maximal compression. Reconstruction accuracies measured by Pearson correlation.

|  | No compression | Maximal compression |
| --- | --- | --- |
| chr1 | 0.64 ± 0.18 | 0.33 ± 0.24 |
| chr2 | 0.65 ± 0.18 | 0.32 ± 0.25 |
| chr3 | 0.67 ± 0.17 | 0.33 ± 0.25 |
| chr4 | 0.64 ± 0.18 | 0.33 ± 0.25 |
| chr5 | 0.64 ± 0.18 | 0.32 ± 0.25 |
| chr6 | 0.65 ± 0.18 | 0.29 ± 0.25 |
| chr7 | 0.64 ± 0.18 | 0.31 ± 0.23 |
| chr8 | 0.65 ± 0.18 | 0.32 ± 0.24 |
| chr9 | 0.65 ± 0.18 | 0.33 ± 0.24 |
| chr10 | 0.64 ± 0.18 | 0.31 ± 0.26 |
| chr11 | 0.66 ± 0.17 | 0.32 ± 0.24 |
| chr12 | 0.65 ± 0.18 | 0.31 ± 0.23 |
| chr13 | 0.65 ± 0.18 | 0.30 ± 0.24 |
| chr14 | 0.65 ± 0.18 | 0.33 ± 0.24 |
| chr15 | 0.65 ± 0.18 | 0.32 ± 0.25 |
| chr16 | 0.64 ± 0.17 | 0.30 ± 0.23 |
| chr17 | 0.67 ± 0.17 | 0.31 ± 0.25 |
| chr18 | 0.68 ± 0.18 | 0.33 ± 0.27 |
| chr19 | 0.65 ± 0.18 | 0.30 ± 0.23 |
| chr20 | 0.66 ± 0.17 | 0.32 ± 0.26 |
| chr21 | 0.69 ± 0.17 | 0.31 ± 0.24 |
| chr22 | 0.66 ± 0.17 | 0.31 ± 0.25 |
| average | 0.65 ± 0.18 | 0.32 ± 0.25 |

**Supplementary Table 3:**
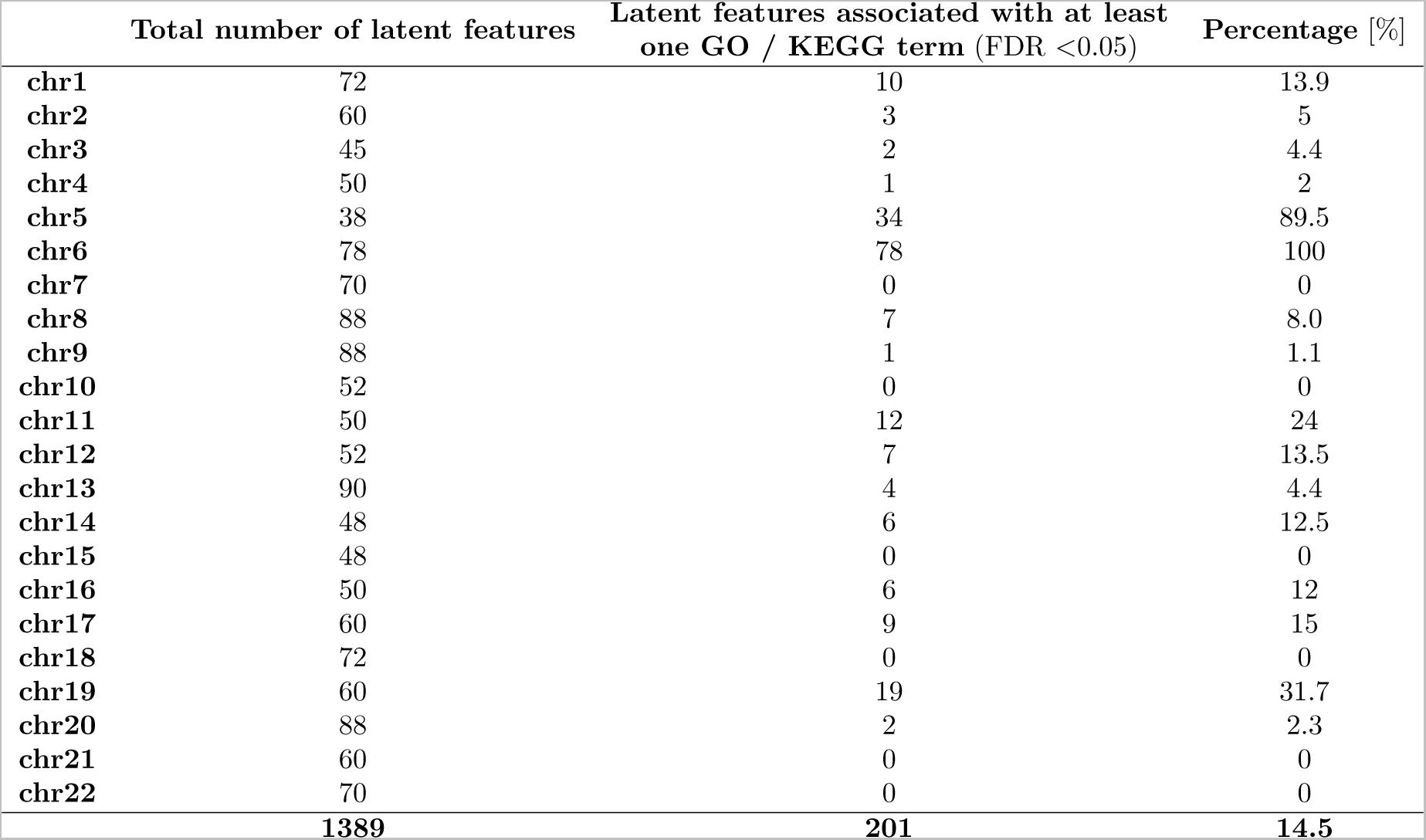
Details on the number of latent features significantly (FDR *<*0.05) associated with GO or KEGG terms per latent feature.

