## Supplementary Material for "mEthAE: an Explainable AutoEncoder for methylation data"

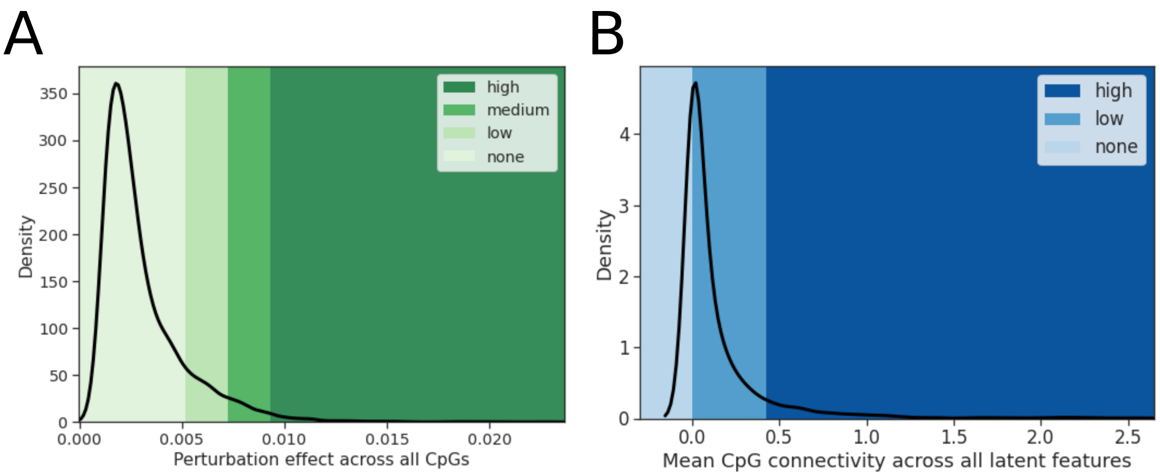


Supplementary Figure 1: Example of observed perturbation effects using chromosome 22;(A) (Local) perturbation effect across all CpGs for a random latent feature. Colours indicate the applied thresholds for cluster assignment, for details see Table 1. (B) (Global) CpG connectivities across all latent features. Colours indicate the applied thresholds for cluster assignment, for details see Table 2

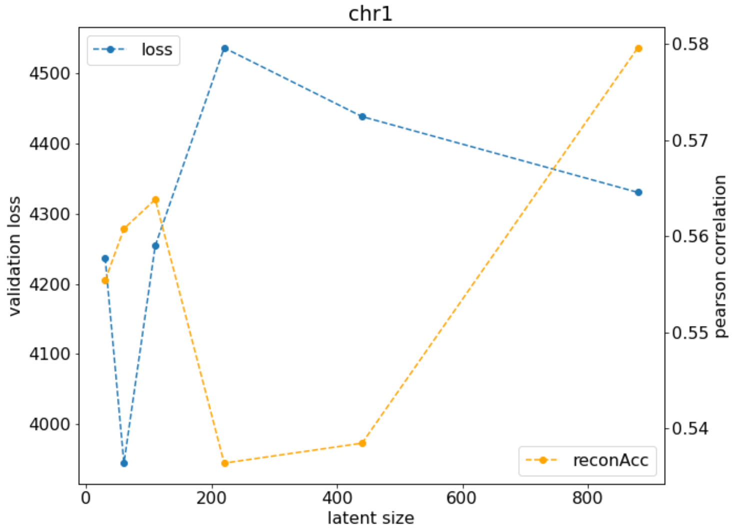


Supplementary Figure 2: Coarse hyperparameter scan results of chromosome 1 (exemplary for other chromosomes); validation loss measured as mean squared error (MSE) loss between input and reconstruction; reconstruction accuracy measured as Pearson correlation between input and reconstruction, averaged over all CpGs

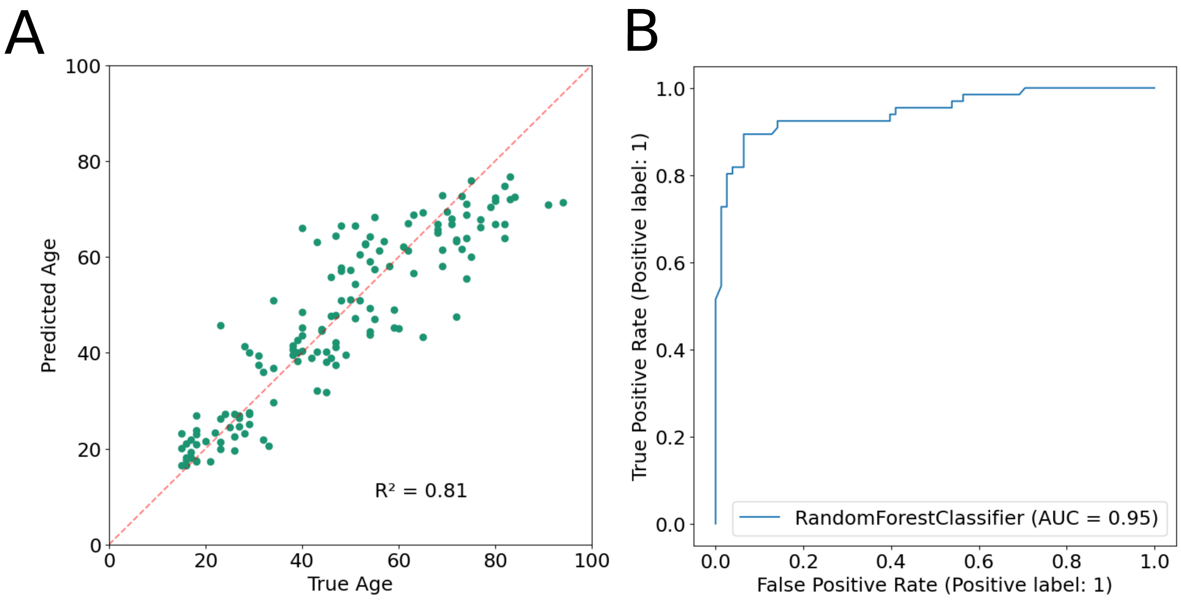


Supplementary Figure 3: Supervised predictions using the combined latent embeddings of all chromosomes, (A) age regression, $R^2$: coefficient of determination; (B) AUC-ROC curve of classification of participants' sex, AUC: Area under the curve

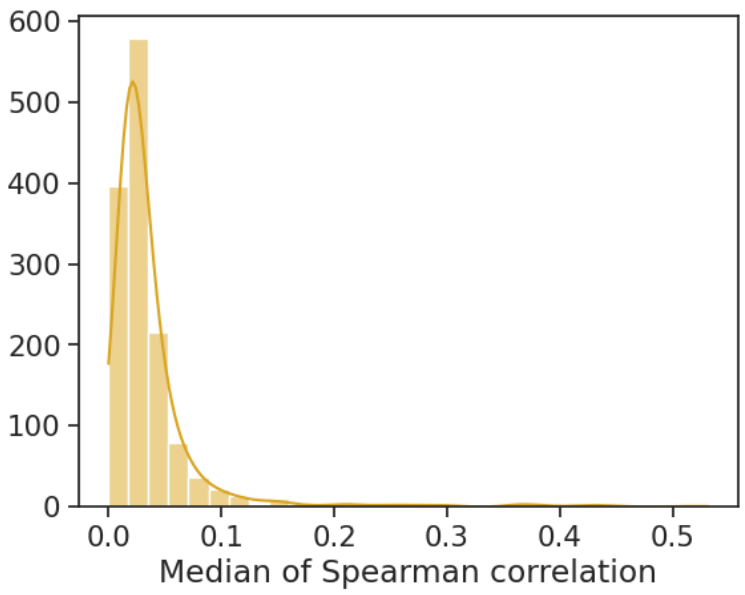


Supplementary Figure 4: Median spearman cross-correlation over all latent features (n=1389)

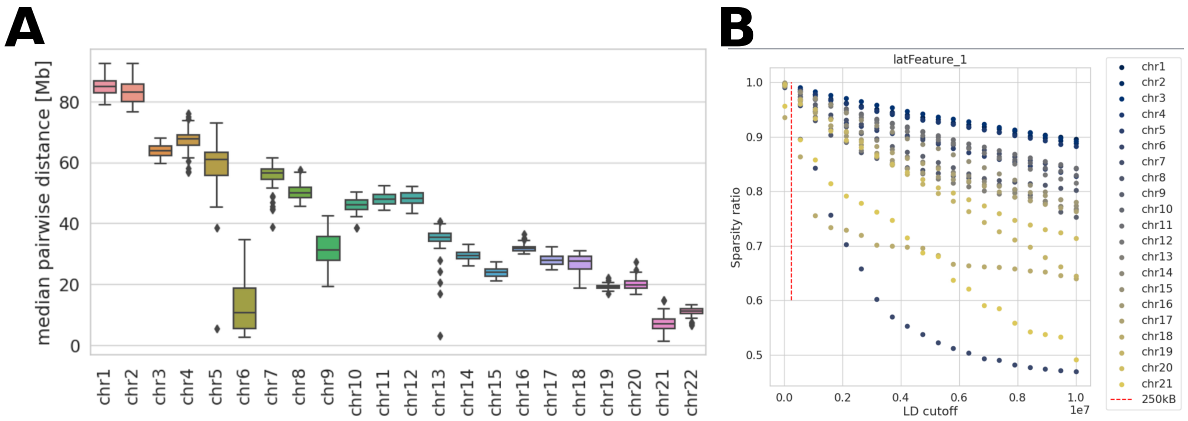


Supplementary Figure 5: (A) Median pairwise distances between local-high CpGs of the same latent feature; all latent features per chromosome shown. (B) Effect of the variation of LD-cutoff values on the sparsity ratio for each chromosome. One latent feature is shown as representative for all latent features.

Supplementary Table 1: Details on architecture of all chromosome-wise autoencoders after hyperparameter optimisation

| | Number of CpGs | Encoder topology | Latent size | Learning rate | Dropout | Compression factor |
|---|---|---|---|---|---|---|
| **chr1** | 29482 | 8840, 1770 | 72 | 0.0001 | 0.1 | 409 |
| **chr2** | 21984 | 6600, 1320 | 60 | 0.0001 | 0.1 | 366 |
| **chr3** | 15547 | 4660, 930 | 45 | 0.0001 | 0.1 | 345 |
| **chr4** | 13100 | 3930, 790 | 50 | 0.0001 | 0.1 | 262 |
| **chr5** | 15965 | 4790, 960 | 38 | 0.0001 | 0.1 | 420 |
| **chr6** | 23070 | 6920, 1380 | 78 | 0.0005 | 0.1 | 296 |
| **chr7** | 19599 | 5880, 1180 | 70 | 0.0001 | 0.1 | 280 |
| **chr8** | 13704 | 4110, 820 | 88 | 0.0005 | 0.3 | 156 |
| **chr9** | 6619 | 1990, 400 | 88 | 0.0005 | 0.3 | 75 |
| **chr10** | 15736 | 4720, 940 | 52 | 0.0001 | 0.1 | 303 |
| **chr11** | 18570 | 5570, 1110 | 50 | 0.0001 | 0.1 | 371 |
| **chr12** | 15165 | 4550, 910 | 52 | 0.0001 | 0.1 | 292 |
| **chr13** | 7802 | 2340, 470 | 90 | 0.0005 | 0.3 | 87 |
| **chr14** | 9417 | 2830, 570 | 48 | 0.0001 | 0.1 | 196 |
| **chr15** | 9715 | 2910, 580 | 48 | 0.0001 | 0.1 | 202 |
| **chr16** | 13823 | 4150, 830 | 50 | 0.0001 | 0.1 | 276 |
| **chr17** | 17044 | 5110, 1020 | 60 | 0.0001 | 0.1 | 284 |
| **chr18** | 3654 | 1100, 220 | 72 | 0.0005 | 0.3 | 51 |
| **chr19** | 15475 | 4640, 930 | 60 | 0.0001 | 0.1 | 258 |
| **chr20** | 6567 | 1970, 390 | 88 | 0.0005 | 0.3 | 75 |
| **chr21** | 2719 | 820, 160 | 60 | 0.0005 | 0.1 | 45 |
| **chr22** | 5243 | 1570, 310 | 70 | 0.0005 | 0.3 | 75 |

Supplementary Table 2: Reconstruction accuracies (measured by Pearson correlation) of models with a latent size equivalent to the number of input CpGs (no compression) and of models with a latent size of 1 (maximal compression)

| | **No compression** | **Maximal compression** |
|---|---|---|
| **chr1** | 0.64 ± 0.18 | 0.33 ± 0.24 |
| **chr2** | 0.65 ± 0.18 | 0.32 ± 0.25 |
| **chr3** | 0.67 ± 0.17 | 0.33 ± 0.25 |
| **chr4** | 0.64 ± 0.18 | 0.33 ± 0.25 |
| **chr5** | 0.64 ± 0.18 | 0.32 ± 0.25 |
| **chr6** | 0.65 ± 0.18 | 0.29 ± 0.25 |
| **chr7** | 0.64 ± 0.18 | 0.31 ± 0.23 |
| **chr8** | 0.65 ± 0.18 | 0.32 ± 0.24 |
| **chr9** | 0.65 ± 0.18 | 0.33 ± 0.24 |
| **chr10** | 0.64 ± 0.18 | 0.31 ± 0.26 |
| **chr11** | 0.66 ± 0.17 | 0.32 ± 0.24 |
| **chr12** | 0.65 ± 0.18 | 0.31 ± 0.23 |
| **chr13** | 0.65 ± 0.18 | 0.30 ± 0.24 |
| **chr14** | 0.65 ± 0.18 | 0.33 ± 0.24 |
| **chr15** | 0.65 ± 0.18 | 0.32 ± 0.25 |
| **chr16** | 0.64 ± 0.17 | 0.30 ± 0.23 |
| **chr17** | 0.67 ± 0.17 | 0.31 ± 0.25 |
| **chr18** | 0.68 ± 0.18 | 0.33 ± 0.27 |
| **chr19** | 0.65 ± 0.18 | 0.30 ± 0.23 |
| **chr20** | 0.66 ± 0.17 | 0.32 ± 0.26 |
| **chr21** | 0.69 ± 0.17 | 0.31 ± 0.24 |
| **chr22** | 0.66 ± 0.17 | 0.31 ± 0.25 |
| average | 0.65 ± 0.18 | 0.32 ± 0.25 |

Supplementary Table 3: Details on the number of latent features significantly (FDR <0.05) associated with GO or KEGG terms per latent feature

| | **Total number of latent features** | **Latent features associated with at least one GO / KEGG term** (FDR <0.05) | **Percentage** [%] |
|---|---|---|---|
| **chr1** | 72 | 10 | 13.9 |
| **chr2** | 60 | 3 | 5 |
| **chr3** | 45 | 2 | 4.4 |
| **chr4** | 50 | 1 | 2 |
| **chr5** | 38 | 34 | 89.5 |
| **chr6** | 78 | 78 | 100 |
| **chr7** | 70 | 0 | 0 |
| **chr8** | 88 | 7 | 8.0 |
| **chr9** | 88 | 1 | 1.1 |
| **chr10** | 52 | 0 | 0 |
| **chr11** | 50 | 12 | 24 |
| **chr12** | 52 | 7 | 13.5 |
| **chr13** | 90 | 4 | 4.4 |
| **chr14** | 48 | 6 | 12.5 |
| **chr15** | 48 | 0 | 0 |
| **chr16** | 50 | 6 | 12 |
| **chr17** | 60 | 9 | 15 |
| **chr18** | 72 | 0 | 0 |
| **chr19** | 60 | 19 | 31.7 |
| **chr20** | 88 | 2 | 2.3 |
| **chr21** | 60 | 0 | 0 |
| **chr22** | 70 | 0 | 0 |
| | **1389** | **201** | **14.5** |
